# Transcriptional analysis of nitrogen fixation in *Paenibacillus durus* during growth in nitrogen-enriched medium

**DOI:** 10.1101/2020.04.13.040188

**Authors:** Mardani Abdul Halim, Quok-Cheong Choo, Amir Hamzah Ahmad Ghazali, Mustafa Fadzil Farid Wajidi, Nazalan Najimudin

**Author notes:** **Corresponding Authors**, Nazalan Najimudin & Mardani Abdul Halim, School of Biological Sciences, Universiti Sains Malaysia, 11800 Penang, Malaysia.

## Abstract

*Paenibacillus durus* strain ATCC 35681^T^ is a Gram-positive diazotroph that displayed capability of fixing nitrogen even in the presence of nitrate or ammonium. However, the nitrogen fixation activity was detected only at day 1 of growth when cultured in liquid nitrogen-enriched medium. The transcripts of all the *nifH* homologues were present throughout the 9-day study. When grown in nitrogen-deficient medium, nitrogenase activities occurred from day 1 until day 6 and the *nifH* transcripts were also present during the course of the study albeit at different levels. In both studies, the absence of nitrogen fixation activity regardless of the presence of the *nifH* transcripts raised the possibility of a post-transcriptional or post-translational regulation of the system. A putative SigA box sequence was found upstream of the transcription start site of *nifB1*, the first gene in the major nitrogen fixation cluster. The upstream region of *nifB2* showed a promoter recognisable by SigE, a sigma factor normally involved in sporulation.

**Significance and Impact of the Study:** *Paenibacillus durus* strain ATCC 35681^T^ is a nitrogen fixing Gram positive bacterium with an unconventional physiological characteristic of being able to fix nitrogen even in the presence of either nitrate or ammonium. It has a total of 6 *nifH* homologues in its genome. In this study, we analysed the transcriptional levels of the *nifH* homologues when grown under nitrogen-enriched and nitrogen-depleted medium. Under nitrogen-enriched condition, the nitrogen fixation activity was detected only at day 1 of growth but the transcripts of all the *nifH* homologues were detected during the course of the study from day 1 until day 9. In nitrogen-deficient condition, nitrogen fixation activities were recorded from day 1 until day 6 and the *nifH* transcripts were present throughout the study. The absence of nitrogen fixation activity even in the presence of the *nifH* transcripts raised the possibility of a post-transcriptional or post-translational regulation of the system.

## Introduction

Nitrogen, the most abundant element on earth, exists in the form of dinitrogen gas and cannot be metabolized by most organisms. However, microorganisms known as diazotrophs are able to convert it to ammonia (Puri et al., 2015). This nitrogen fixation process requires an enzyme called nitrogenase (Dixon and Kahn, 2004). Most regulatory studies on nitrogen fixation are based on the Gram-negative model using *Klebsiella pneumoniae* and *Azotobacter vinelandii* (Dixon, 2004, Kennedy and Bishop, 2004). *Clostridium pasteurianum* was one of the few Gram-positive diazotrophs used as a model. It has six *nifH* homologs and their upstream regions contained a consensus sequence different from the usual NifA box of *K. pneumoniae* (Wang et al., 1988, Johnson et al., 1993). NifA serves as a transcriptional regulator in *K. pneumoniae* that binds to the NifA box. However, this gene is absent in *C. pasteurianum* indicating that the regulation of the *nif* genes is probably different (Chen, 2004).

Members from the genus *Paenibacillus* are now emerging as models for studying nitrogen fixation in Gram positive bacteria. Genomic analysis showed that 9 core genes are involved nitrogen fixation are arranged as a cluster containing *nifB*, *nifH*, *nifD*, *nifK*, *nifE*, *nifN*, *nifX, hesA* and *nifV* (Xie et al., 2014). The *Paenibacillus* sp. WLY78 cluster produced an active nitrogenase when expressed *Escherichia coli* using a σ^A^-dependent promoter upstream of *nifB* (Wang et al., 2013).

*Paenibacillus durus* ATCC 35681 has the ability to undergo nitrogen fixation even in the presence of fixed-nitrogen (Seldin et al., 1984a). An acetylene reduction analysis revealed that the nitrogenase enzymes was not inhibited by the presence of nitrate (Seldin et al., 1984a). Subsequent analysis showed that the expression of its nitrogenase genes occurred even in the presence of high levels of ammonium (Teixeira et al., 2008). The strain showed the presence of multiple *nifH* homologues (Choo et al., 2003). A whole genome analysis of ATCC 35681 revealed six copies of *nifH* including those coding for the alternative nitrogenases (Halim et al., 2016). The presence of multiple copies of *nifH* generates an interest to understand their expression profiles, especially under nitrogen-enriched and nitrogen-limited conditions. Here, we report an expression study of the six *nifH* homologues as well as measurements of nitrogenase activity of ATCC 35681 cells grown under both conditions. The transcription start sites (TSS) of the *nifH* and *nifB* genes were also determined to assess the presence of potential promoters in their upstream regions.

## Results and discussion

### Nitrogenase activity of ATCC 35681 in nitrogen-limited and nitrogen-enriched media

A diauxic growth pattern was obtained when ATCC 35681 was cultured in nitrogen-limited medium (Fig. S1). Nitrogenase activity was detected as early as day 1 in the nitrogen-limited medium reaching a peak at the beginning of the first lag phase on day 3. The reading subsequently decreased and no activity was observed from day 7 onwards (Fig. 1).

**Fig. 1.**
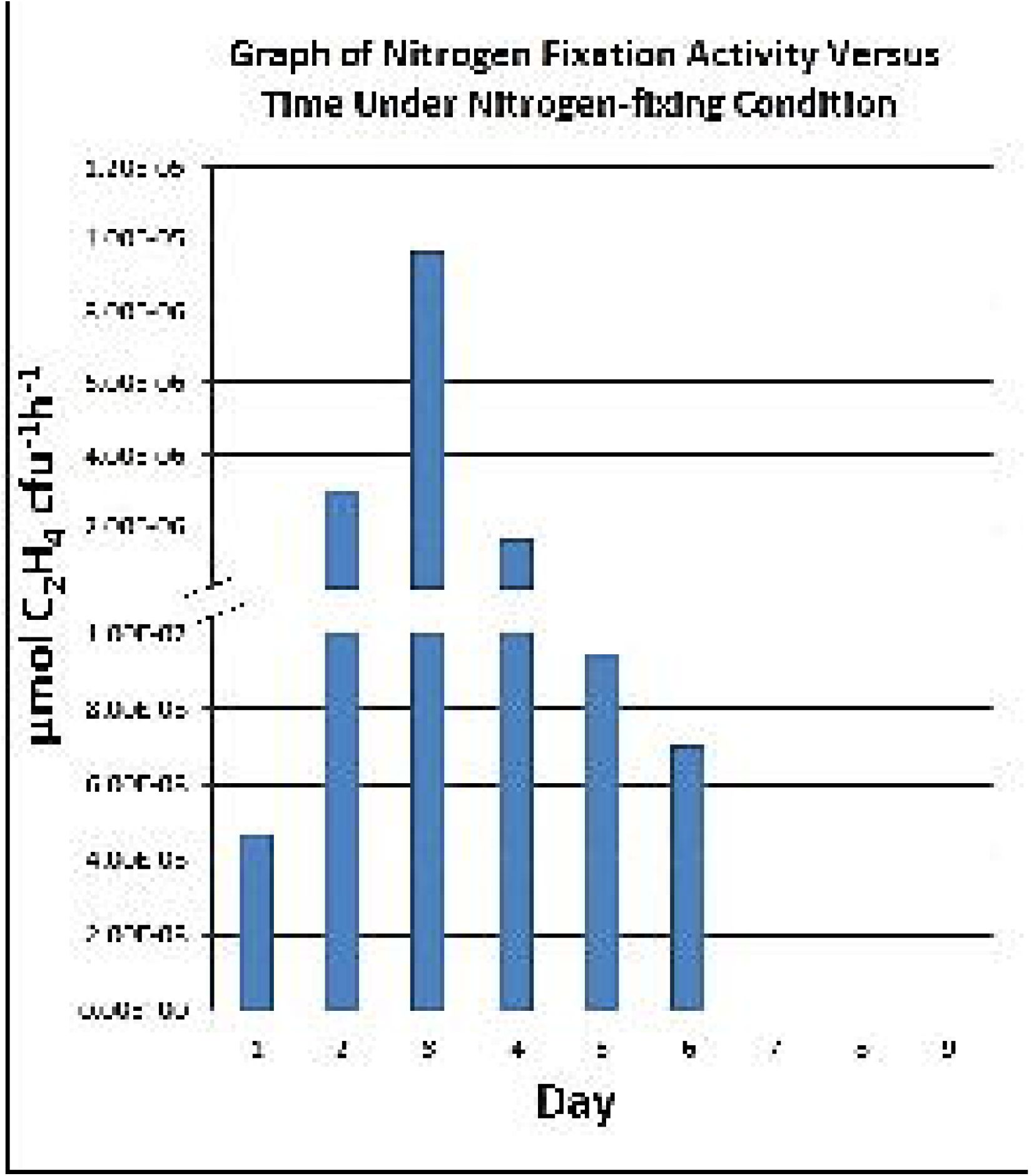
Graph of ATCC35681 nitrogenase activity versus time under nitrogen-limited (nitrogen-fixing) condition. Nitrogenase activity was detected as early as day 1 in the nitrogen-limited medium reaching a peak at the beginning of the first lag phase on day 3. The reading subsequently decreased and no activity was observed from day 7.

In nitrogen-enriched medium, the typical sigmoidal growth curve was obtained with the mid-exponential stage reached at day 2 and the stationary stage at day 4. Nitrogen fixation activity was detected on day 1 and this ability to fix nitrogen even in the presence of ammonium was unusual. No activity was observed from day 2 until day 9 although the cells were viable. The lag phase is normally a period of slow growth during which the cells are adapting to their new environment and synthesising cellular components using pre-assembled macromolecules (Rolfe et al., 2012). It may include reparation of damages accumulated during prior growth. ATCC 35681 is able to perform nitrogen fixation in the presence of fixed nitrogen in the early lag phase possibly by using the pre-assembled nitrogenase enzyme produced during their prior stage of growth. Free-living diazotrophic bacteria are generally subjected to immediate inhibition of nitrogen fixation activity or repression of nitrogenase synthesis in response to external ammonium or nitrate ions (Hartmann et al., 1986, Streeter and Wong, 1988). They respond to the fluctuating environmental conditions by regulating the enormously demanding nitrogen fixation process. During high availability of external fixed nitrogen, most diazotrophs regulate nitrogen fixation at the transcriptional level. Species related to ATCC 35681 such as *Paenibacillus macerans* and *Paenibacillus polymyxa* also showed total inhibition of nitrogen fixation activity in the presence of 0.5 % nitrate (Seldin et al., 1984a).

The ability to fix nitrogen in the presence of nitrate during a 24-hour growth period described by (Seldin et al., 1984a) was investigated further using qRT-PCR. In this study, nitrogen fixation did occur during growth in nitrogen-enriched condition but only in day 1. Interestingly, transcripts of the *nifH* genes were measurable during the course of growth showing that all the genes were expressed albeit at different levels. Under nitrogen-enriched condition, the genes *nifH2, nifH4, nifH5* and *nifH6* showed maximum expression at the early stage of growth. The highest expression was shown by *nifH4* and this was observed in day 1 with an NRQ value of 3.94. All the genes were generally displayed their lowest expression during the late stationary stage on day 8 and 9.

These observations raised the possibility of ATCC 35681 having a post-transcriptional mode of regulation to prevent nitrogen fixation from occurring after day 1. A possible mechanism of post-transcriptional regulation is via small RNA molecules (sRNAs). Bacteria have evolved sRNAs to adjust gene expressions in response to changes in the environment.

These are non-coding RNA molecules of about 50-500 nucleotides long and are highly organized containing s everal stem and loop structures (Waters and Storz, 2009). In a root-associated bacterium, *Pseudomonas stutzeri* A1501, a novel regulatory non-coding RNA known as NfiS adjusts the subsequent total nitrogenase activity by directly pairing with the *nifK* mRNA and thus regulating its translational efficiency. The interaction also affects the half-life of the mRNA (Zhan et al., 2016). Small RNAs are also known to work in conjunction with the proteins Hfq and CsrA in controlling the expressions of numerous regulons. Hfq acts as a chaperone by simultaneously binding to both sRNA and its target mRNA to enable a successful base-pairing interaction (Geissmann and Touati, 2004, de Almeida Ribeiro Jr et al., 2012). In contrast, CsrA directly binds to the 5’ regions of mRNAs and represses protein synthesis (Liu and Romeo, 1997, Schubert et al., 2007).

Another form of post-transcriptional regulation has been discovered in the nitrogen fixation system of the filamentous cyanobacterium *Anabaena variabilis* ATCC 2941 in which nitrogen fixation occurred only during the early stage of growth. It has three nitrogenase genes. Two of these code for Mo-nitrogenase enzymes which are present in which one was heterocyst-specific while the other functions only under anoxic conditions in vegetative cells and heterocysts (Thiel, 2004). The third enzyme is a V-nitrogenase that is also heterocyst specific. Only one type of nitrogenase is functional under different physiological conditions and these genes are post-transcriptionally regulated (Thiel and Pratte, 2014). Transcripts were initiated upstream of its *nifHDK1* and *vnfH* genes, and these were subsequently truncated to generate 5’-ends which developed secondary structures that affected the stability of the transcripts (Ungerer et al., 2010). The resultant differential expression ensured that the cells made the correct enzyme for the environmental condition in which it was growing in.

In the nitrogen-fixing medium, the expressions of each *nifH* gene was elevated higher compared to the nitrogen-enriched during the course of the diauxic growth. Nitrogen fixation activity was also observed during day 1 to day 6. The genes *nifH2* and *nifH3* were expressed extraordinarily higher during the late stationary period of growth than the lag and exponential phases of the second diauxic growth. This suggested that their expressions were developmentally linked to the growth cycle. Spores were observed at this stage of growth coinciding with the period that nitrogen fixation was occurring. The second cycle of the diauxic growth showed a relatively reduced nitrogen fixation activity. The cells concomitantly advanced to a more pronounced sporulation stage. The nitrogen fixation activity continued during the second cycle of diauxic growth until the late stationary stage during which most cells had already sporulated. The expression of all six *nifH* genes from day 1 until day 9 for both under nitrogen-enriched and nitrogen-limited conditions, respectively, are presented in (Fig. S2)

In prokaryotes, sigma factors of the RNA polymerase holoenzyme recognized distinct sets of promoters to initiate transcription. The transcription start site (TSS) specified by the 5’end of a transcript provides important information about promoters and the associated sigma factors. The major cluster of genes containing *nifB1H1D1K1ENX*-*hesA*-*nifV* was investigated. The TSS for *nifB1* revealed no discernible promoter in the upstream region. The normally conserved −35 and −10 regions of TTGACA and TATAAT of the housekeeping SigA-bound promoters were absent. However, two regions bearing the sequence TGT-N10-ACA were uncovered at a more distal positions of −60 and −72 from the TSS (Fig. 2). These regions resembled the NifA boxes which are specifically recognized by the NifA protein in various proteobacterial diazotrophs (Buck et al., 1986, Jacobson et al., 1989, Fischer, 1994, Lee et al., 2000). In spite of this, a search in the genome of ATCC 35681 failed to reveal a *nifA* homologue. The absence of a conventional bacterial promoter motif also raised the possibility of the involvement of *trans-acting* factors.

**Fig. 2.**
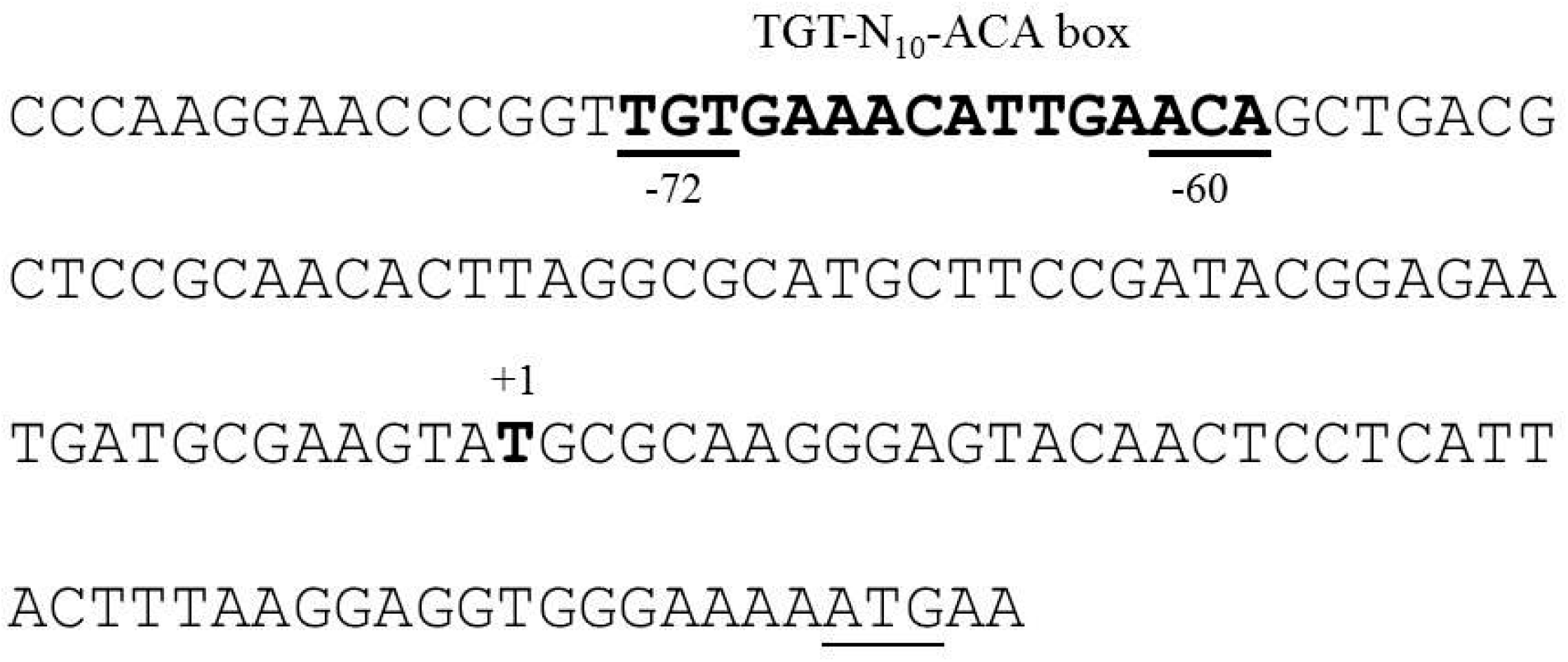
Putative promoter of *nifB1* gene. The underlined ATG represent start codon for *nifB1*. The bold (+1) T represent putative transcription start site. Putative promoter located −60 and −72 preceding +1. −60/−72 region resemble NifA box.

In the intergenic region between *nifB1* and *nifH1*, a TSS was unexpectedly found and −12 region of TATTAATA highly resembled a consensus promoter sequence (TTTA(T/A)ATA) of the *nif* genes of methanogenic *Methanococcus maripaludis* and several other archaea (Reeve, 1993). The −45 region (TTGAAA) closely mirrored the −35 promoter region of Gram-positive bacterial SigD (TTGACA) (Wang et al., 1988) (Fig. 3). This *nifH1* promoter could possibly be active only under certain physiological or environmental conditions. The presence of internal promoters has been identified in several polycistronic operons of bacterial gene clusters. For example, the histidine biosynthesis operons of *E. coli* and *Salmonella enterica* consist of eight polycistronic genes and two internal promoters were recognised for the genes *hisC* and *hisF*. (Carlomagno et al., 1988). The expression of such promoters depends on the physiological conditions that the cells are encountering (Grisolia et al., 1983).

**Fig. 3.**
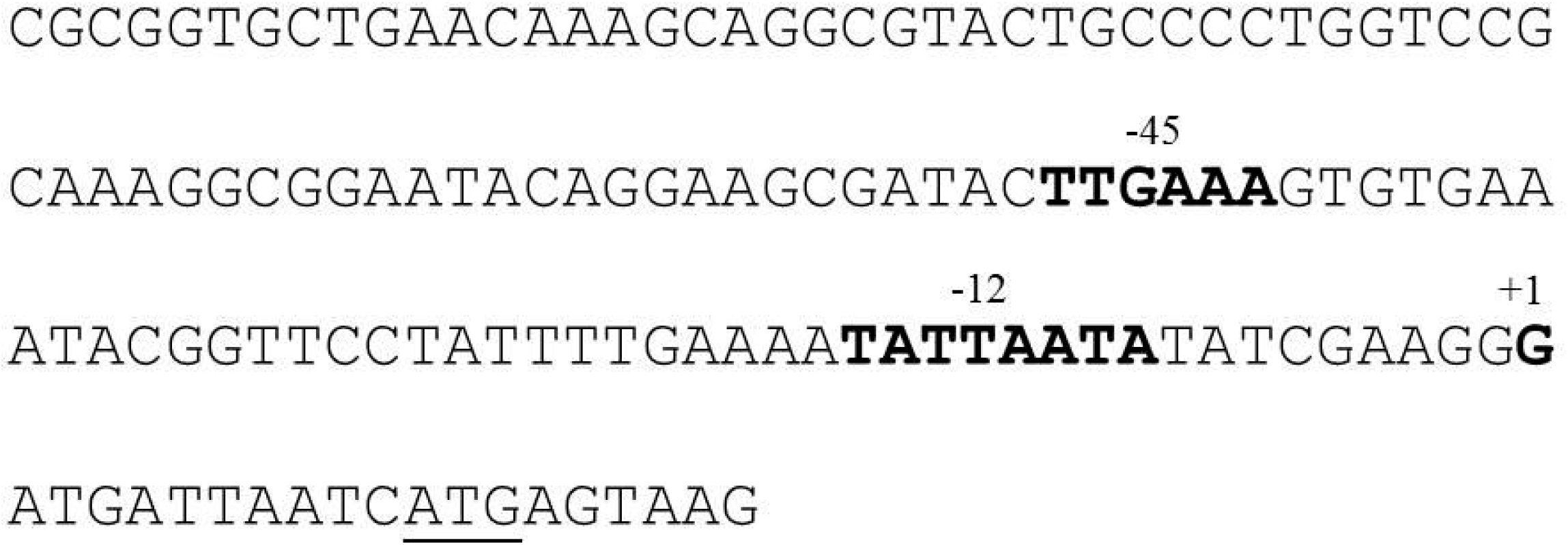
Putative promoter of *nifH1* gene. The underlined ATG represent start codon for *nifH1*. The bold (+1) G represent putative transcription start site. Putative promoter located − 12 and −45 preceding the transcription start site. The −12 region of *nifH1* highly resembled a consensus promoter sequence (TTTA(T/A)ATA) of the *nif* genes of methanogenic bacteria and from several archaea. The −45 region closely resembled the −35 promoter region of Gram-positive bacterial SigD.

The gene *nifB2* is located as an operon with *nifH2*. Its TSS was mapped to a T-residue, 21 bp preceding its putative ATG translational start site (Fig. 4). The predicted −75 and −45 promoter sequences of TAAGATG and CATACGAA, respectively, shared considerable homology with the *Bacillus subtilis* SigE-regulated promoter sequences of (T/G)(A/C)ATA(A/T)(A/T) and CATACA(A/C)T (Roels et al., 1992). SigE is a sigma factor coded by *spoIIG* and it plays an important role during the process of sporulation by creating a polar division of the dividing cell (Haldenwang, 1995). The functional sigma factor is formed by a proteolytic activation of a precursor protein into a mature form that directs the expression of SigE-dependent genes in the mother cell compartment. The presence of a SigE-like promoter sequences upstream of the *nifB2-nifH2* operon indicated that this sigma factor is possibly engaged in the control of the expression of the *nifB2-nifH2* operon during this stage of growth.

**Fig. 4.**
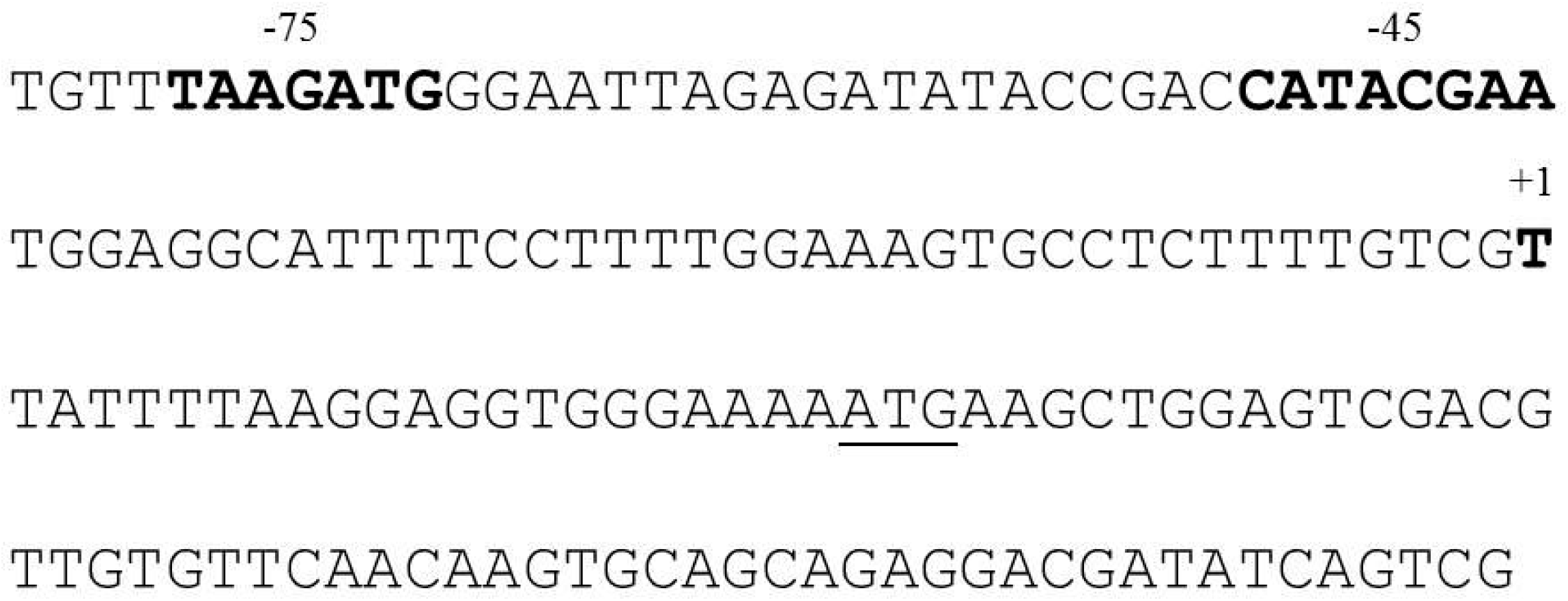
Putative promoter of *nifB2* gene. The underlined ATG represent start codon for *nifH1*. The bold (+1) T represent putative transcription start site. The putative promoter located −45 and −75 preceding the transcription start site. The predicted promoter region resembled *Bacillus subtilis* SigE-regulated promoter sequences

The absence of a distinguishable consensus SigN-type promoter upstream of the *nifH* genes indicated that different are being recognised by SigN of ATCC 35681. A similar situation exist in *Clostridium pasteurianum* in which none of the promoter regions of its six *nifH* homologs possessed a consensus sequence recognised by SigN (Wang et al., 1988). Moreover, two of the *C. pasteurianum nifH* genes had sequences similar to the consensus −35 and −10 promoter regions of *E. coli* and to the Gram-positive promoter recognised by SigD of *B. subtilis*. In the latter, the SigD factor is needed for the transcription of the flagellin and motility genes necessary for chemotaxis (Chen et al., 2009). SigD plays also an essential role in the regulation of autolysins that hydrolyse and remodel the peptidoglycan found in the bacterial cell wall.

## Materials and methods

### Bacterial strain, plasmids and growth conditions

*Paenibacillus durus* ATCC 35681 was obtained from the American Type Culture Collection. The strain was maintained aerobically with continuous shaking in GB (glucose broth) medium at 30 °C (Seldin et al., 1984b). Thiamine-biotin-nitrogen medium (TBN) was used for growth involving nitrogen-fixing condition as previously described (Seldin and Penido, 1986). TBN has a limited nitrogen source in the form of yeast extract to initiate growth. For growth under nitrogen-enriched condition, 4 mM of NH_4_Cl was added to TBN (Teixeira et al., 2008). Plasmid vector pGEMT^®^-T (Promega, USA) was used to clone the amplified DNA fragments containing the 5’ends of cDNA. *Escherichia coli* JM109 (*sup*E44, *end*A1, *hsdR17, gyrA96, rel*A1, *thi*Δ(*lac-pro*AB), *rec*A1, F’[*tra*D36*pro*AB^+^*lac*I^q^*lac*ZΔM15] (courtesy of SA Zahler, Cornell University, USA) was used in this study as the host strain for plasmids.

### Growth curve determination

The growth curves of ATCC35681 were plotted from day 1 until day 9 for both nitrogen-enriched and nitrogen-limited media.

### Acetylene reduction assay (ARA)

Nitrogenase activities were determined using ARA as previously described (Bergersen, 1970). ARA was performed daily with the final reading expressed as μmol C_2_H_4_ cfu^-1^ h^-1^. All experiments were performed in triplicate.

### Total RNA isolation

Total RNA samples were isolated using TRIzol reagent as described by the manufacturer’s protocol (Invitrogen, USA). The extracted RNA was purified with RNeasy^®^ MinElute^®^ Cleanup Kit (Qiagen, Germany) and subsequently treated with RNase-free DNase (Qiagen, Germany) to remove any trace of genomic DNA. The integrity of RNA was verified by agarose gel electrophoresis and the concentration was determined by using Nanodrop 2000 (Thermoscientific, USA).

### Quantitative real-time PCR (qRT-PCR) analysis

All six *nifH* genes were chosen for the qRT-PCR analysis. Total RNA was extracted from cells grown in both nitrogen-enriched (TBN plus NH4Cl) and nitrogen-limited (TBN) conditions from day 1 until day 9. The iTaq^™^ Universal SYBR^®^ Green One-Step Kit (Bio-Rad, USA) was used to perform the qRT-PCR. All the experiments were performed in triplicate. The three housekeeping genes used as internal control were *gapd*, *gyrb* and *pyrk*. All the primers used for qRT-PCR are listed in Table S1.

### Determination of 5’ ends of mRNA

The 5’-ends of the mRNA molecules of *nifB* and *nifH* genes were determined as described in the manual of GeneRacerTM RNA ligase-mediated rapid amplification of cDNA ends (RLM-RACE) kit (Invitrogen, USA) which were based on previously established methods (Kazuo and Sumio, 1994, Volloch et al., 1994, Schaefer, 1995). A sum of 10 μg of total RNA was ligated to the synthetic RNA oligonucleotide adapter (5’-CGACUGGAGCACGAGGACACUGACAUGGACUGAAGGAGUAGAAA-3’). The ligated mRNA was reverse-transcribed into cDNA using the SuperScript^™^ III RT kit (Invitrogen, USA). All the primers used to reverse-transcribe and to amplify the junctional 5’-fragment of the cDNA are listed in Table S2. The PCR products generated from the amplification were gel purified, cloned into pCR^®^4-TOPO^®^ vector (Invitrogen, USA) and subsequently transformed into *E. coli* JM109 competent cells. Positive clones were subjected to DNA sequencing to determine the 5’-ends of the *nifH* and *nifB* genes.

## Acknowledgements

We gratefully acknowledge all of our collaborators and colleagues for the discussion and the work conducted in this lab. We would like to thank the funding received from Universiti Sains Malaysia under under the Research University Program (1000/PBIOLOGI/811184).

